# The Genome Explorer Genome Browser

**DOI:** 10.1101/2024.04.24.590985

**Authors:** James Herson, Markus Krummenacker, Aaron Spaulding, Paul O’Maille, Peter D. Karp

## Abstract

Are two adjacent genes in the same operon? What is the order and spacing between several transcription-factor binding sites? Genome browsers are software data-visualization and exploration tools that enable biologists to answer questions such as these. In this paper we report on a major update to our browser, Genome Explorer, that provides nearly instantaneous scaling and traversing of a genome, enabling users to quickly and easily zoom into an area of interest. The user can rapidly move between scales that depict the entire genome, individual genes, and the sequence; Genome Explorer presents the most relevant detail and context for each scale. By downloading the data for the entire genome to the user’s web browser and dynamically generating visualizations locally, we enable fine control of zoom and pan functions and real-time redrawing of the visualization, resulting in smoother and more intuitive exploration of a genome than is possible with other browsers. Further, genome features are presented together, in-line, using familiar graphical depictions. In contrast, many other browsers depict genome features using data tracks, which have low information density and can visually obscure the relative positions of features. Genome Explorer diagrams have high information density that provides larger amounts of genome context and sequence information to be presented in a given sized monitor than for tracks-based browsers. Genome Explorer provides optional data tracks for analysis of large-scale datasets and a unique comparative mode that aligns genomes at orthologous genes with synchronized zooming.

## 1 Importance

Genome browsers provide graphical depictions of genome information to speed uptake of complex genome data by scientists. They provide search operations to help scientists find information, and zoom operations to enable scientists to view genome features at different resolutions. We introduce the Genome Explorer browser which provides extremely fast zooming and panning of genome visualizations, and displays with high information density.

## 2 Introduction

Genome browsers communicate the positions of functional elements within a genome to scientists, and support inference of new genome features from large datasets. These functional elements include genes, transcription start sites, transcription-factor binding sites, and origins of replication. Genome browser designers also hope to enable efficient navigation through a genome that will enable scientists to interpret experimental datasets with respect to genome organization, compare related genomes, and extract and export genome-sequence regions.

In more detail, the problems that genome browser designers seek to solve include the following. In order to effectively convey the full range of features and spatial relationships within a genome, browsers must be able to scale their graphical presentations from the sequence level to a level where an entire prokaryotic chromosome is displayed in one screen, a factor of approximately 1500 (from 10 bases per inch to approximately 15KB per inch). This scaling must be done quickly and smoothly to enable the user to rapidly find the scale that answers their current informational question.

At these many scales, browser designers face the problem of conveying an appropriate information density [1] (meaning the screen area required to display a given piece of information) that enables scientists to find the information they want, as well as providing surrounding genome context, without forcing the user to endlessly engage with zoom and positional controls (which can be quite slow for older browsers if the server must generate a new image for every such change). Another challenge browser designers face is to provide useful semantic zooming levels. Semantic zooming successively reveals new graphical features at different zoom levels, such as gene names and transcription start sites.

The Pathway Tools genome browser has been under development since 1995 [2, 3, 4]. This article describes its third incarnation, which we call Genome Explorer. Genome Explorer is notable for employing a different graphical organization than most genome browsers, which are predominately organized around a series of parallel visual “tracks.” Although Genome Explorer does support tracks, it is primarily organized around genome diagrams that capture genome features in a manner that is both more space efficient than tracks, and that communicates spatial relationships, including superposition, more effectively than do tracks.

Many genome browsers have been implemented over the years and have made use of a number of computer technologies. Early, first-generation browsers were desktop-based, including AceDB [5] and the first incarnation of the Pathway Tools genome browser [2, 3]. The development of the World Wide Web in the 1990s led to second-generation browsers that used image-based web technologies including GBrowse [6, 7], the Ensembl genome browser [8], the NCBI genome browser [9], the IMG genome browser [10, 11], the MicroScope genome browser [12], and the second incarnation of the Pathway Tools genome browser [4]. Second-generation browsers are relatively slow because their genome images are generated on a remote server and each zoom operation generates a new image that must be downloaded from the server via the internet, which can take a second or more.

The third generation of faster web-based genome browsers use JavaScript to generate the genome images within the user’s web browser, and include JBrowse [13], JBrowse 2 [14], newer versions of the UCSC Genome Browser [15], and Genome Explorer. Although third-generation genome browsers are certainly faster than second-generation browsers, there is still significant variation among their capabilities. Here we present the capabilities of Genome Explorer.^1^

## 3 Results

Genome Explorer can operate in three different modes: basic mode supports search and browsing of a single replicon; comparative mode supports comparison of two or more genomes aligned at orthologous genes; and tracks mode enables visual analysis of large-scale datasets such as chip-seq data.

Genome Explorer is part of the Pathway Tools software, which powers the BioCyc.org website and a number of other websites. Genome Explorer is available for use with all of the 20,000 genomes within BioCyc.org, each of which is stored in a Pathway/Genome Database (PGDB). Experiment with the browser at this URL with the free EcoCyc database for *E. coli* K–12: https://biocyc.org/genbro/genbro.shtml?orgid=ECOLI&replicon=COLI-K12. Different BioCyc databases vary as to which genome features they contain, such as transcription-start sites and terminators; therefore, different sets of features will be visible in the genome browser for different databases. EcoCyc has a particularly comprehensive collection of information.

### 3.1 Basic Browsing Mode

An example Genome Explorer window is presented in Figure 1. This window depicts the basic mode of Genome Explorer. In basic mode the major components of the Genome Explorer window are indicated by numbered regions in the diagram as follows. (1) Genome Explorer search bar. (2) Command buttons (orange) and zoom-level selectors (blue). (3) Depiction of full length of current replicon; the red rectangle indicates the region shown at higher resolution below. (4) Highresolution area of replicon. (5) Legend explaining graphical conventions used in high-resolution area. The legend is invoked at user request. The check-boxes in the legend enable and disable display of each type of feature in the high-resolution area. The On and Off settings within the legend are absolute; under the Auto setting visibility of a feature is computed by semantic zooming rules.

**Figure 1.**
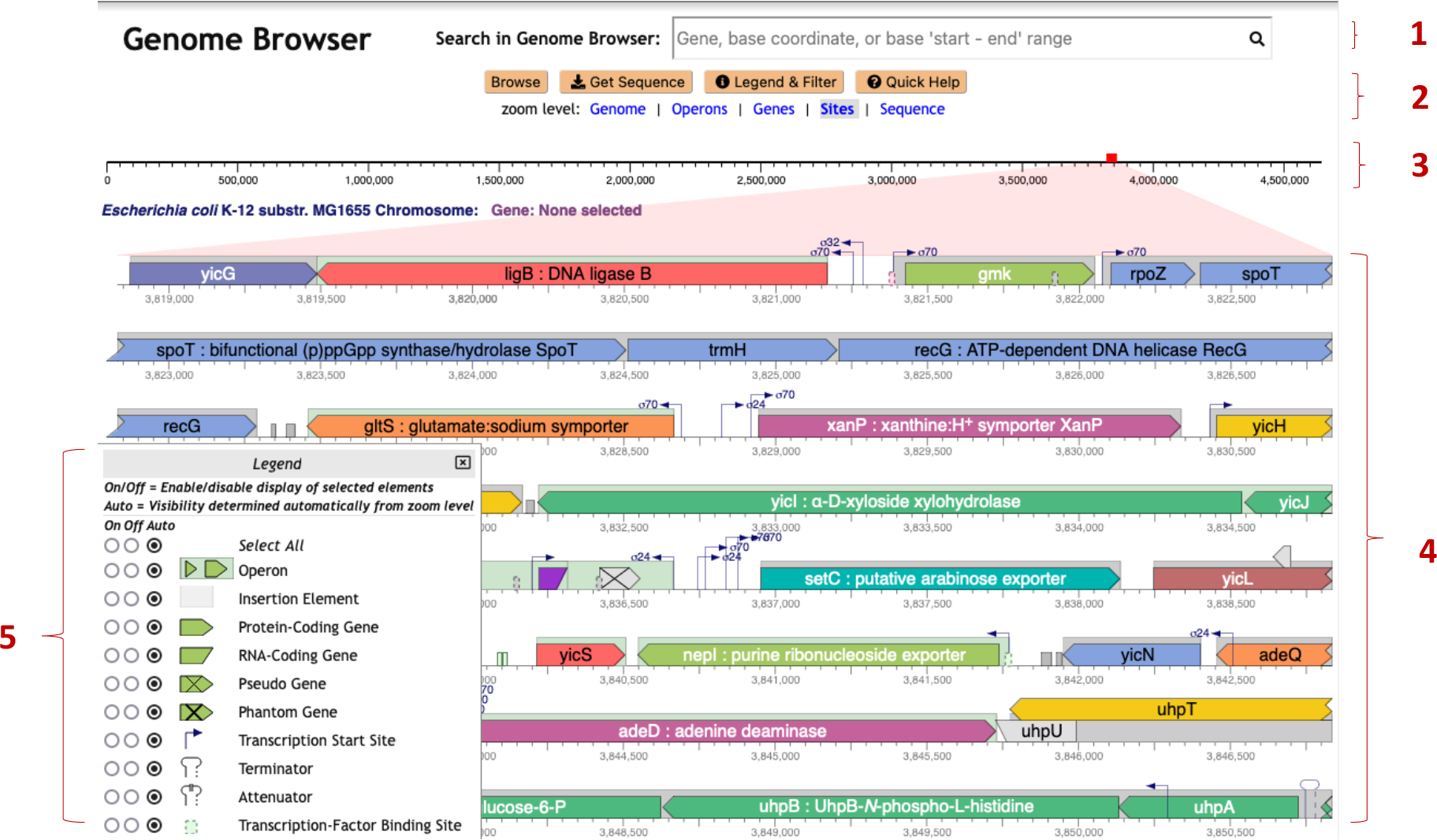
Genome Explorer basic mode, with legend shown in the lower-left corner. Numbers are explained in the text.

Within the high-resolution area (4) multiple graphical icons depicting genes and genome sites are shown. Lines wrap vertically as do the lines of a book. Gene color indicates operon organization: adjacent genes in the same color belong to the same operon. The gray boxes indicate the extents of operons. As indicated by the legend, this image depicts protein-coding genes (example: *ligB* in the top line), RNA-coding genes (example: short purple gene to the right of the legend), and pseudogenes (example: gray gene with an “X” to the right of the purple RNA-coding gene). The Genome Explorer does not yet depict introns and exons, which are planned for future work, hence currently the Genome Explorer is best suited for bacterial genomes. A variety of sites are shown here including transcription start sites (with sigma-factor indicated), terminators (last line), and transcription-factor binding sites (examples: two green sites to the right of the legend and to the left of *yicS*).

Even within this small window shown for publication purposes, a fairly large region of the genome encompassing many operons is shown because of line wrapping, yet there is also room to depict fairly small sites such as transcription-start sites. We refer to the display of genes, transcription-start sites, terminators, and other sites adjacent to one another within the same rectangular regions as “in-line display.”

#### 3.1.1 Navigation: Zooming, Translation, and Search

Genome Explorer zooming operations are performed by spinning the mouse wheel, scrolling the trackpad, or pressing the up/down arrow keys, while pointing the mouse at the desired centerpoint for the zooming operation (such as the upstream region of a gene). In this fashion the user can ensure the area they point at remains on the screen for the duration of the zooming operation.

As we increase the zoom level around a given region using the Genome Explorer, more and more information becomes visible. Gene names and product names are depicted as the size of each gene increases. Transcription factor names appear (see Figure 2, first line) as do the names of binding sites for small RNAs. The legend (see Figure 1) depicts the full set of genome features that are depicted. Further zooming reveals the nucleotide sequence and the amino-acid sequence of coding regions (Figure 3). Zooming out reveals overall genome organization (Figure 4). As shown in that figure, tooltips are available at all zoom levels to provide additional information on genes and sites.

**Figure 2.**
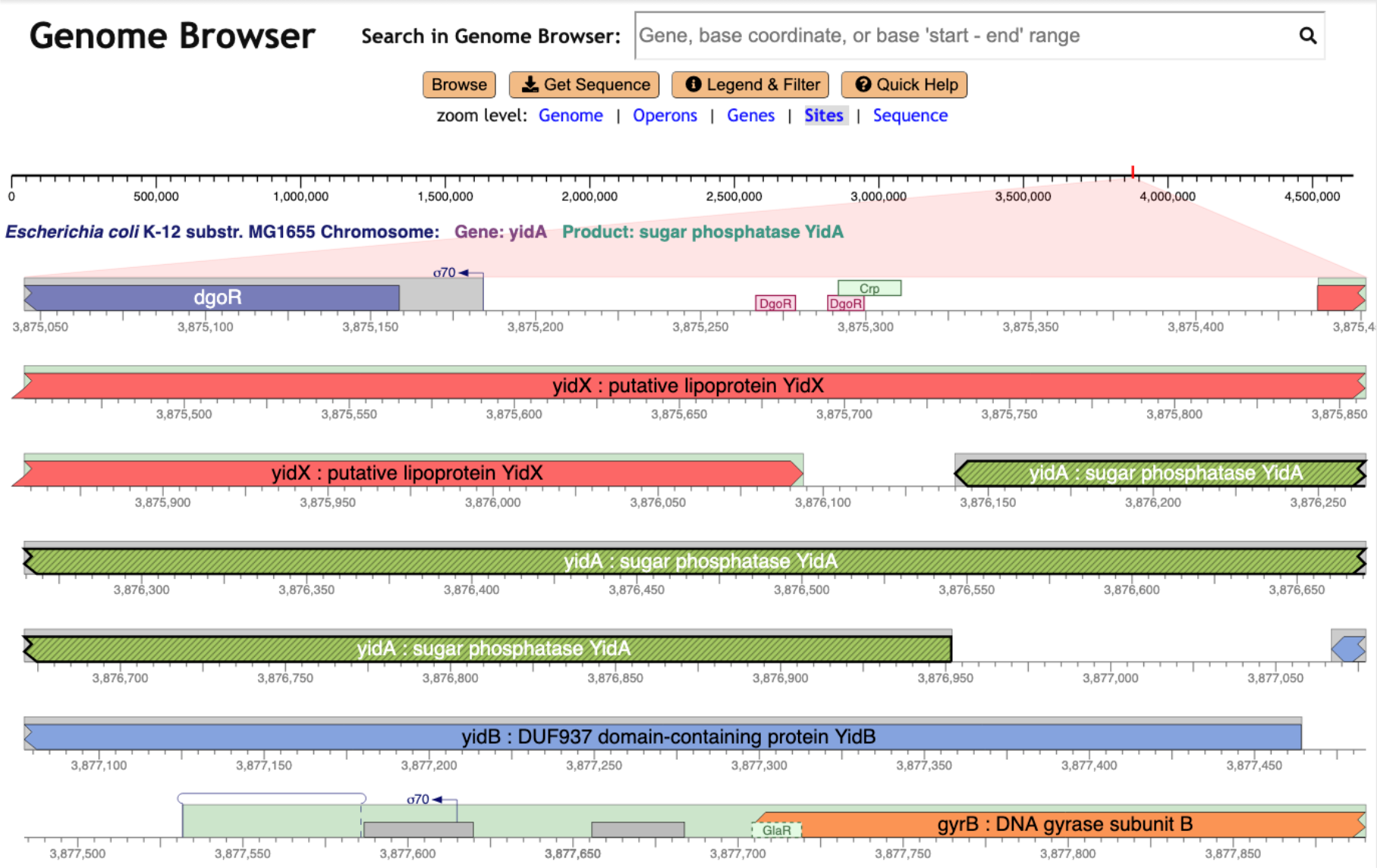
Genome Explorer zoomed to depict sites in the *E. coli* genome, after a search for the *yidA* gene, which is highlighted.

**Figure 3.**
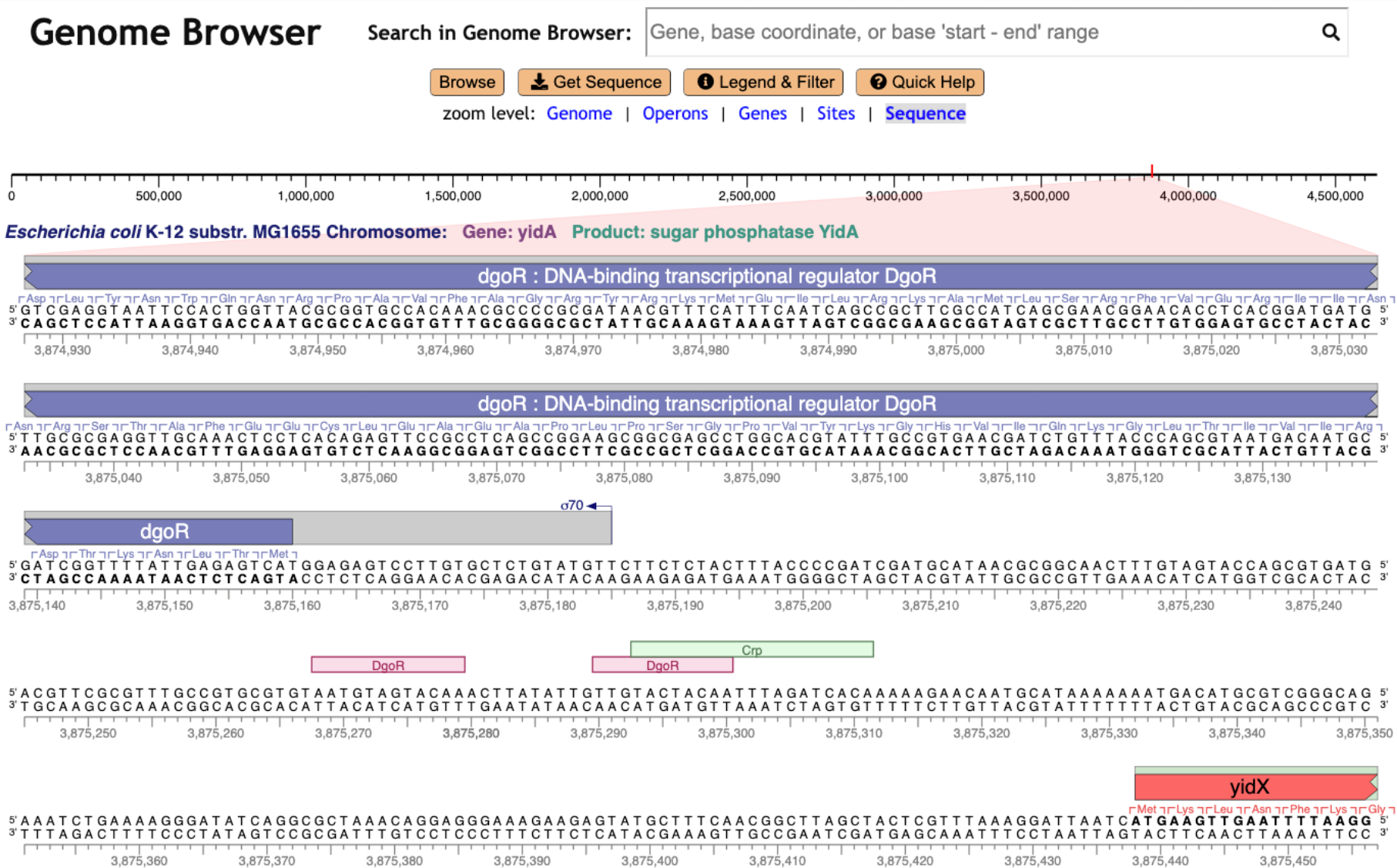
Genome Explorer zoomed to depict sequence in the *E. coli* genome.

**Figure 4.**
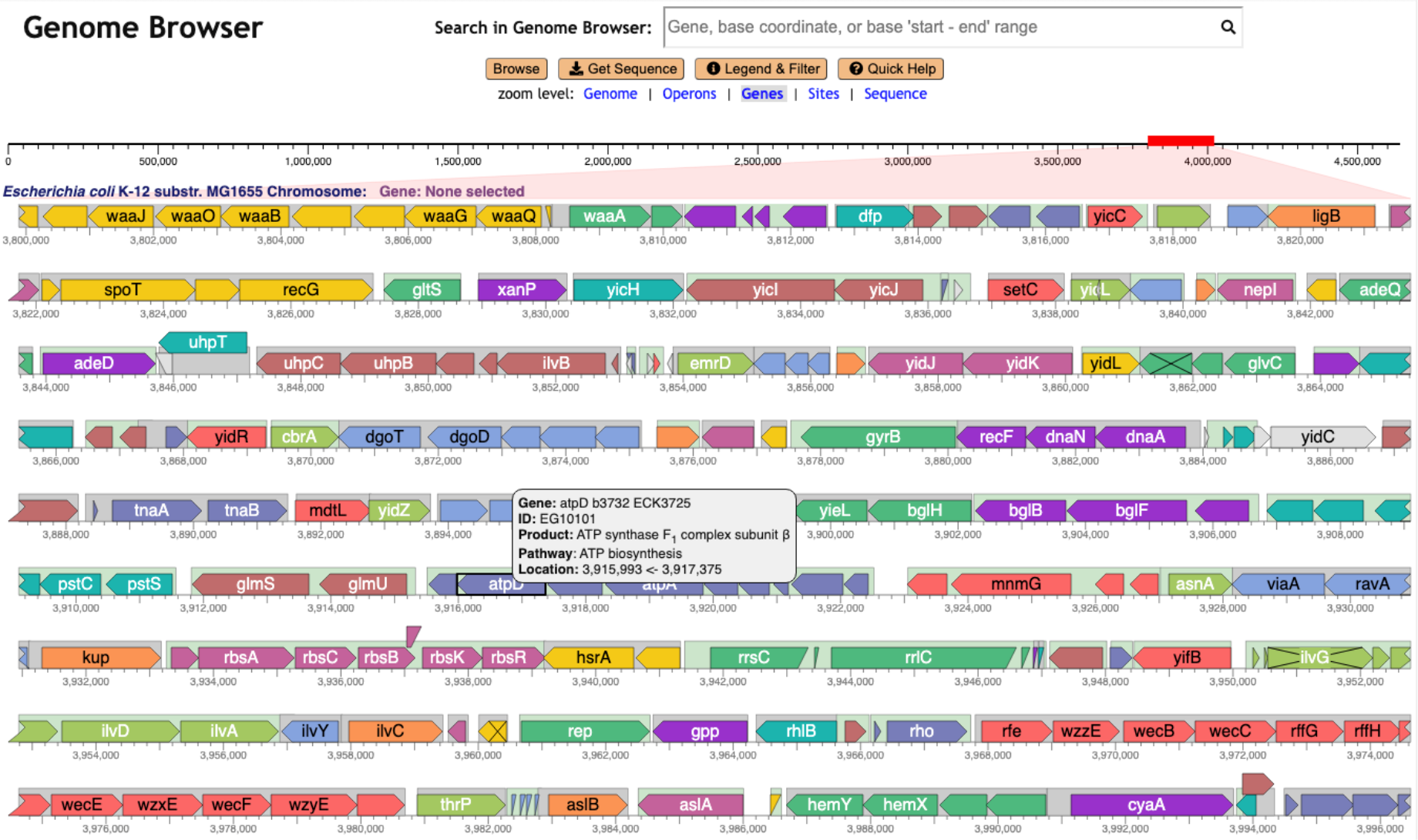
Genome Explorer displays a 200kb region of the *E. coli* genome.

Zooming can also be performed by clicking on the zoom levels listed under component (2) in Figure 1; for example, clicking on “Sequence” zooms immediately to the sequence level.

The user can move horizontally within the genome by clicking and dragging with the mouse, such as by dragging a gene left, right, up, or down. The user can also move horizontally by dragging the red box in the full-replicon diagram at the top, by clicking at a position within that diagram, and by pressing the left-arrow and right-arrow keys.

The “Search in Genome Browser” box shown at the top of Figure 1 and several other figures can be used to position the browser at a feature of interest based on a user-supplied gene name, accession number, gene-product name (including substrings), single base coordinate, or start/end coordinates.

The use of the mouse wheel and trackpad provide fine control over the amount of zooming that occurs. In contrast, click-based zooming occurs at rather coarse increments that can be quite difficult to adjust to achieve the exact desired scaling – coarser scaling improves zooming speed but increases the difficulty of arriving at exactly the desired zoom level.

#### 3.1.2 Selection of Nucleotide and Amino-Acid Sequences

Basic mode provides a sequence-selection capability whereby the user zooms to the starting base (or amino-acid residue) of interest, clicks on it, and then zooms to the ending base (or residue) and clicks on that. There is no limit to the size of the selected region, and for circular chromosomes the selected region can span the origin. The selected nucleotide or amino-acid sequence region can be copied to the clipboard or saved to a FASTA file.

Other browsers supporting sequence selection include IMG, NCBI, JBrowse, and the UCSC browser.

### 3.2 Comparative Mode

Our goal in developing the comparative mode of Genome Explorer is to enable users to easily visualize differences in the conservation of genes and other features across many genomes. Figure 5 shows an example comparison across several strains of *E. coli* that can be re-created using the URL https://biocyc.org/genbro/ortho.shtml?lead-orgid=ECOLI&lead-genes=EG11024&orgids=GCF_001021615,ECOLI,GCF_000010765,GCF_004010715,GCF_900636075. Instead of using sequence-based alignments, comparative mode aligns genomes at orthologous genes. The user invokes comparative mode by specifying a “lead gene” in a given organism, and a set of other organisms to compare with. Genome Explorer includes in the alignment all of the user-selected organisms that have an ortholog to the lead gene, based on the ortholog database maintained by BioCyc. The genomes are aligned at the center-point of each ortholog. Each replicon is drawn in one line — line wrapping is disabled in comparative mode.

**Figure 5.**
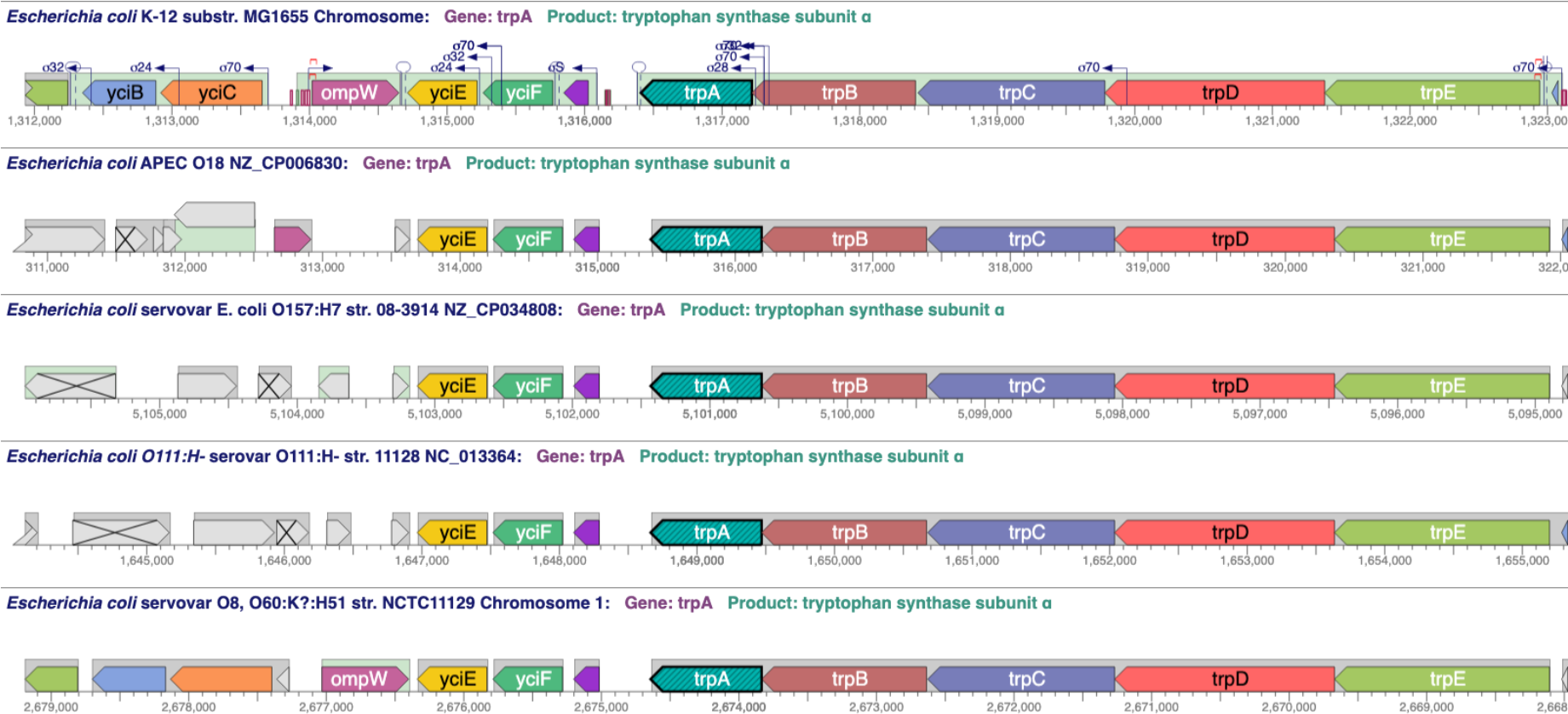
Genome Explorer comparative mode applied to the region around the trp operon in five *E. coli* strains. Genes in color have an ortholog in the top strain whereas gray genes have no ortholog in the top strain.

The meaning of the gene colors is different in comparative mode: genes in the same colors are orthologs, but with the caveat that only a dozen colors are available, and colors are recycled after the dozen have been used, so some genes in the same color are not orthologs. However, usually it is clear from gene position, name, and length, which genes are orthologs and which are not. To be completely sure, the user can hover the mouse over a given gene, which visually highlights all of its visible orthologs.

Comparative mode depicts all other genome features present in the displayed region for each genome. Zooming and panning are controlled in the same way as for basic mode; the genomes zoom and pan in a synchronized fashion. The user can select a different lead gene at any time.

We are not aware of other browsers that support an ortholog-based comparative mode or that provides synchronized panning and zooming.

### 3.3 Tracks Mode

Tracks mode enables the analysis of one or more large-scale datasets visually aligned against the genome to correlate features in those datasets with known genome features such as genes that are stored in the PGDB. When tracks are enabled the Genome Explorer changes to an unwrapped (single-line) in-line display and the one or more input datasets are drawn below that single-line display (see Figure 6). Zooming and panning of the tracks region and of the in-line diagram are synchronized and use the same mouse gestures as does basic mode.

**Figure 6.**
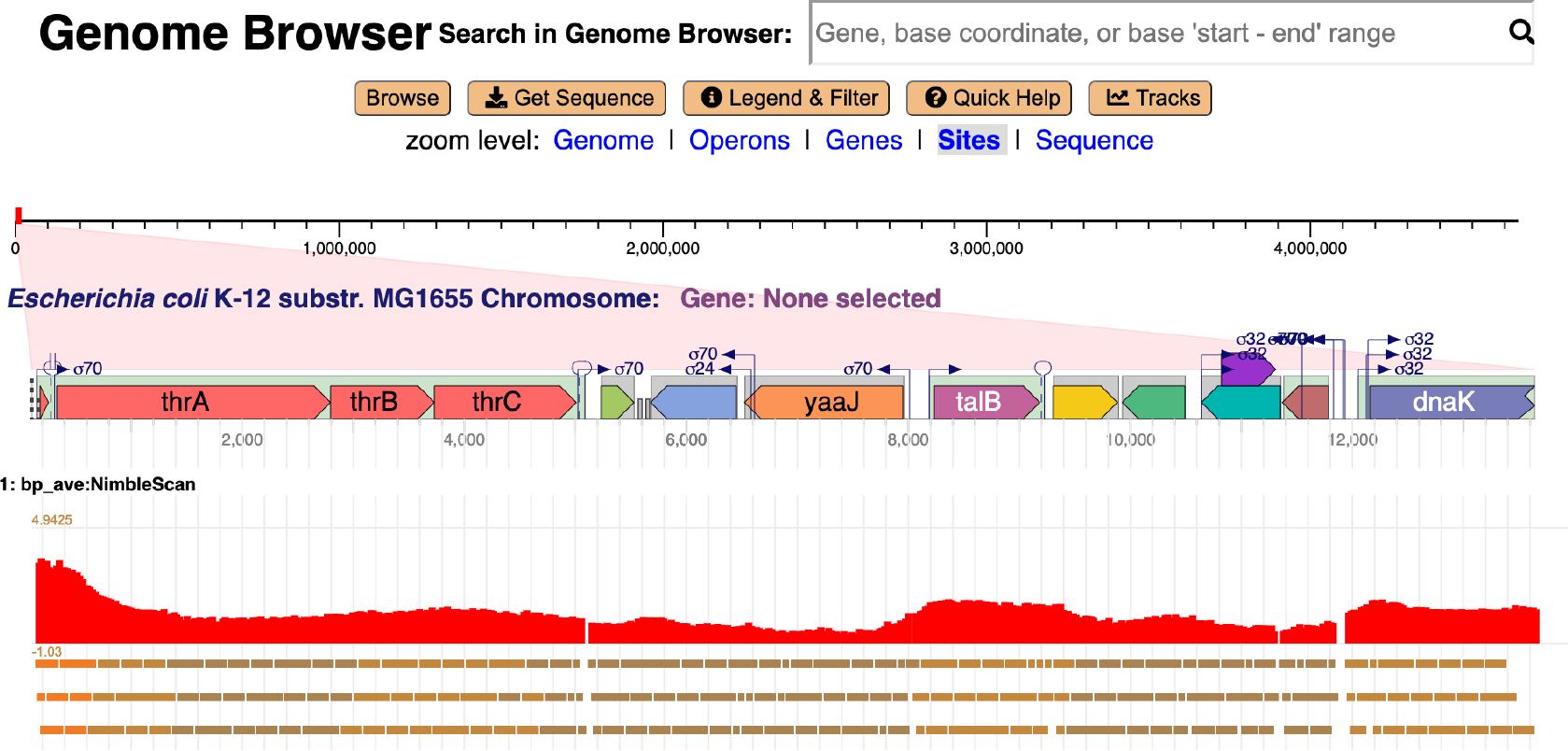
Genome Explorer tracks showing intensity of RNA polymerase binding in a section of the *E. coli* genome. The same dataset is represented twice in the diagram: the three linear regions at the bottom of the figure show binding regions as rectangles; the color of each rectangle indicates the intensity of binding. The red bar graph just above the three linear regions depicts the intensity of binding using the Y-axis.

Track data can be drawn in three different styles, two of which are shown in Figure 6. Track data can be drawn as horizontal bars that indicate the genomic extent of each feature in the track data file. The color of each bar reflects the intensity value, if any, provided in the input data file for that genome region. Track data can also be drawn as a bar graph (red graph in Figure 6) or point graph (not shown) for cases in which the input data include an intensity value for the Y-axis. A tracks control panel (not shown) enables the user to select the display style and Y-axis scale for each track. The Y-axis scale is needed because the scale of the data can vary greatly in different regions of the genome, thus the default scale from the minimum to the maximum data value is not appropriate for every region of the genome.

The Genome Explorer accepts tracks data in the GFF file format. The data shown in Figure 6 is available at http://www.ai.sri.com/pkarp/pubs/genome-explorer-tracks.gff.

## 4 Discussion

### 4.1 Tracks Compared with In-Line Display

Browsers such as JBrowse, GBrowse, and the UCSC browser make extensive use of data tracks in the sense that tracks are the primary visual mechanism for representing every type of genome feature. For example, the JBrowse window in Figure 7 provides, in downwards order, tracks for operons (red), transcription start sites, terminators, and ribosome binding sites. Each site is shown as a small rectangle with a direction-indicating arrow. In Genome Explorer the preceding types of information are displayed in-line alongside the gene diagrams. One advantage of the in-line approach is that it is more efficient in its use of vertical space, enabling Genome Explorer to wrap multiple lines and display much more of the genome within a screen of a given size, while still depicting many types of sites.

**Figure 7.**
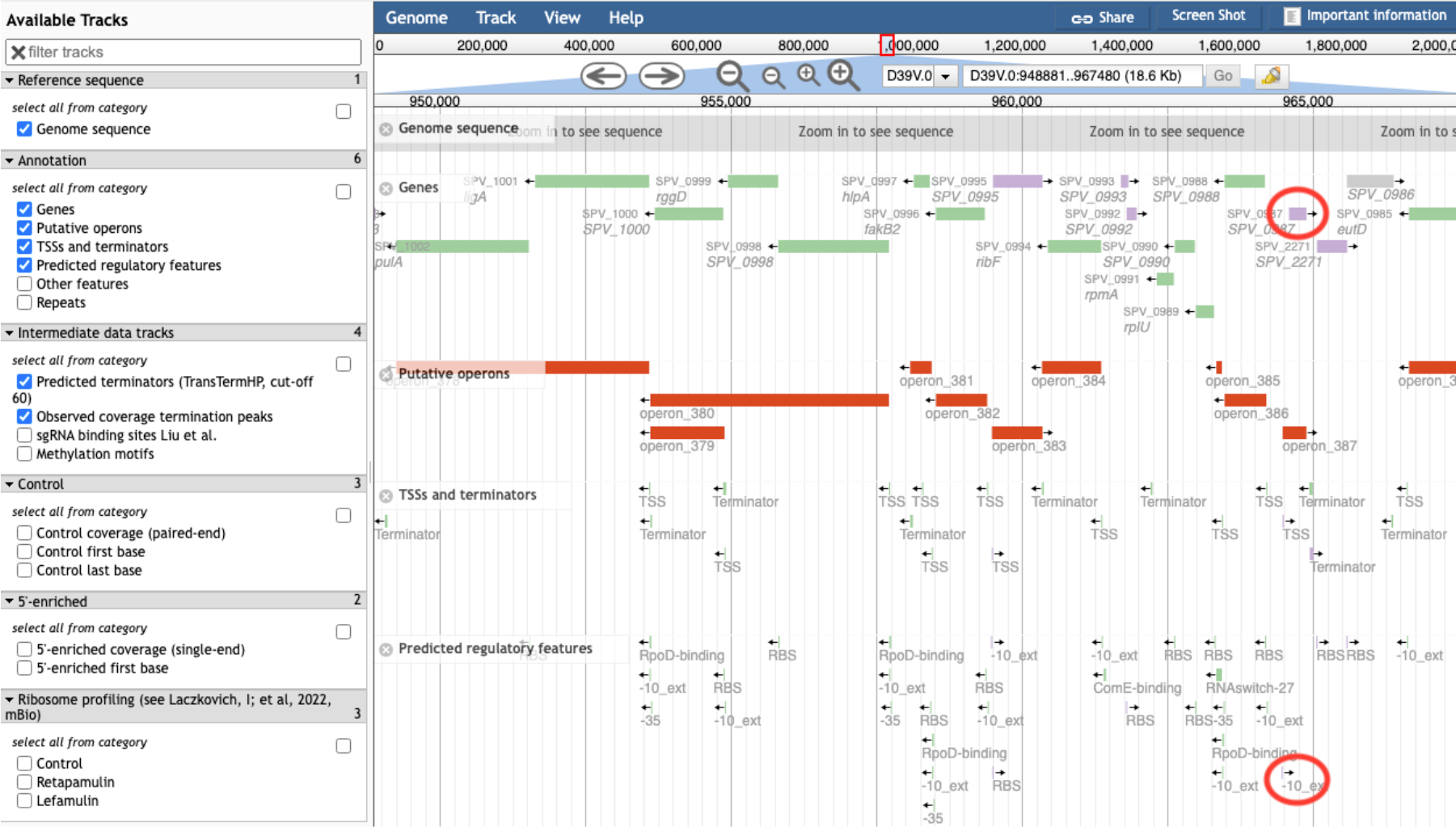
A JBrowse view of the *Streptococcus pneumoniae* D39V genome. Several tracks are present in this view; for example, the green and purple rectangles near the top constitute the “Genes” track, and the red rectangle constitutes the “Putative operons” track.

Compare Figure 1 with a JBrowse window for *Streptococcus pneumoniae* as shown in Figure 7. These windows contain similar numbers of genes, but few gene names and no product names are shown in the JBrowse window due to insufficient space — because no wrapping is performed. To understand what genes are present in the JBrowse window the user must manually hover over every gene to see its tooltip. Thus, it is more time consuming to extract the same information from a JBrowse page versus a Genome Explorer page.

Higher information density means more of the surrounding genome context is visible, and when the sequence is visible, it means more sequence information can fit in the same size screen. For example, on the same large monitor JBrowse can depict one line containing 125 bases, whereas Genome Explorer can depict 10 wrapped lines containing 2400 bases.

Another issue with using tracks versus in-line display is that the vertical separation of elements on different tracks increases the difficulty in ascertaining the positions of features relative to one another. For example, in Figure 7 it is visually challenging to assess the relative locations of features that are close to one another horizontally but far from one another vertically. Do the two genome features that we have circled in red on the right side of the diagram overlap or not? To answer this question the user must carefully track their eyes vertically from one feature to another and try to measure the distance of each feature to the nearest vertical line, a process that is both time consuming and prone to error. This issue is a fundamental problem with the tracks approach.

In contrast, in a Genome Explorer in-line display, these features are right next to each other and it is trivial and instantaneous to evaluate their relative positions. Overlapping features do present challenges that we often handle through stacking of genes (see Figures 1 and 4), transcription start sites, and transcription factor binding sites (see Figures 2 and 3). At times we simply draw overlapping features on top of one another.

In-line display is also more intuitive to biologists than are tracks, because in-line display uses graphical conventions (e.g., transcription start sites are depicted by arrows) that biologists are familiar with from articles and textbooks, whereas tracks are less familiar.

All this said, tracks are clearly useful and important, particularly for organisms with a large number of diverse experimental datasets that simply cannot all be moved in-line, as occurs when there is no graphical convention for depicting that type of data, or there are too many types of overlapping data in the same horizontal region. For example, the UCSC genome browser provides large numbers of tracks for *Homo sapiens* data. However, the more data that can be moved in-line to reduce the number of tracks shown, the more we simplify the evaluation of positional relationships for those tracks that remain by decreasing the average vertical distance between tracks. Most microbes have many fewer experimental datasets than are available for humans and hence have much less need for large numbers of tracks. Thus, the Genome Explorer use of a hybrid inline and tracks display exploits the strengths of both approaches.

### 4.2 Genome Browser Zooming

We consider rapid, efficient zoom and pan to be key tools for helping users explore and understand a genome. We have optimized these operations to make them as fast and easy as possible. Compared to second-generation browsers, Genome Explorer zooming is very rapid because all of its zooming is computed within the users’s web browser and does not require network communication with the server — thus zooming occurs essentially instantaneously. Browsers that use older web technologies must request the server generate a new image each time a zoom click occurs, and wait for that image to be transmitted across the internet. Compared to other third-generation browsers, we have prioritized zooming over scrolling by re-purposing the mouse wheel and two-finger trackpad swipe for zoom instead of scroll. This approach is also used in other interfaces in which zooming is a key activity such as in maps and many image editors. This is in contrast to other browsers that require clicking a widget. Additionally, the fast response of Genome Explorer permits a “continuous zoom” so that the genome smoothly expands or contracts by small increments around the mouse cursor rather than larger discrete steps. This enables the user to stay better oriented. Finally, our implementation allows the user to easily switch between zoom and pan operations, which are both used to navigate to a desired view –— users don’t have to move the mouse to different areas of the screen for each activity.

Typically, click-based zooming in browsers such as JBrowse provides four zooming buttons: two that zoom in and two that zoom out, with each pair providing a large zoom step and a small zoom step (see the four magnifying glasses near the top of Figure 7). One reason wheel-based zooming is faster is that the user controls the zoom increment by the speed at which they rotate the wheel, whereas with zoom buttons the increments are fixed and are often the “wrong” size for what the user is trying to accomplish, with manual entry of coordinates the only way to interpolate between the provided sizes.

The second reason wheel-based zooming is faster is that when using click-based zooming across very large scales is because it is easy to lose track of one’s position within the genome since most browsers zoom in and out with respect to a fixed point, e.g., the center of the diagram. Often the center of the diagram is not the point the user wants to zoom in on. After clicking a few times, the user becomes lost, having zoomed in to an unfamiliar area of the genome, and can have difficulty figuring out how to get to the region they wanted to go to. The user must spend time orienting themselves and backtracking to earlier in the zooming process, where they can recognize some landmark. In contrast, Genome Explorer zooming uses the mouse pointer position as the fixed point, around which zooming is centered, and thus the user controls the zoom point. With practice one learns to make subtle adjustments to the zoom point as the mouse wheel spins, does not become lost during zooming, and has no reason to backtrack.

## 5 Materials and Methods

Genome Explorer is implemented in JavaScript and uses an HTML5 canvas. It has been tested on Chrome, Firefox, and Safari.

When the user invokes the Genome Explorer on a new genome, the browser makes several Web service calls back to a Pathway Tools server. Those calls return all genome features on the selected replicon, and, for comparative mode, the orthologs among the selected genomes. These services are implemented in Common Lisp.

The speed comes from the fact that all graphics operations are performed in the user’s web browser. The only data retrieved during operation of the browser are chunks of DNA sequence that are requested on demand for the region being drawn.

## Acknowledgments

We thank Suzanne Paley, Peter Midford, and Lisa Moore for helpful suggestions. This work was supported by grant NSF2109898 from the National Science Foundation.

## Footnotes

1 Although the EcoCyc database is free to all users, the site does require users to create a free account after a number of page views.

